# *De novo* design of binder proteins targeting *Helicobacter pylori* adhesin BabA

**DOI:** 10.64898/2026.05.24.727452

**Authors:** Yaojun Zhu, Murtala bindawa isah, Xiaoying Zhang

## Abstract

*Helicobacter pylori* has been classified as a Group 1 carcinogen by the International Agency for Research on Cancer of the World Health Organization and is one of the most well-established risk factors for gastric cancer. Long-term colonization by *H. pylori* depends on adhesin-mediated attachment to the gastric mucosa, among which the blood group antigen-binding adhesin BabA is a key surface factor involved in host recognition, tissue tropism, and persistent infection. In this study, we established a structure-guided computational design pipeline to develop compact protein binders targeting functionally relevant epitopes of BabA. First, using experimentally resolved BabA–antibody and BabA–nanobody complex structures as templates, we extracted structural contact residues on BabA through heavy-atom contact analysis, thereby defining antibody-recognition epitopes supported by complex-structure evidence. In addition, sequence-based, structure-based, and evolutionary conservation analyses were integrated to identify candidate functional epitope residues with high antigenicity, strong conservation, and surface-exposed features. On this basis, constrained *de novo* backbone generation was performed around the prioritized epitope regions, followed by amino acid sequence design and structural back-validation of the candidate binders. Candidate BabA–binder complexes were further evaluated using molecular docking, molecular dynamics simulations, and residue-level interface perturbation analysis to assess interface stability, epitope occupancy, and potential binding hotspots. This workflow enables systematic screening of BabA-targeting binders that may compete with antibody-recognized functional surfaces. Although these candidates still require experimental validation, this study provides a transferable computational framework for designing compact protein binders against pathogen adhesins by integrating experimentally resolved complex-structure resources with computational epitope prioritization based on sequence, conformation, and evolutionary conservation, and establishes a preliminary library of BabA candidate binders for subsequent validation and optimization.

## 1 Introduction

*Helicobacter pylori* is a highly adapted gastric pathogen that primarily colonizes the human stomach and contributes to gastroduodenal diseases, including chronic gastritis, peptic ulcer disease, duodenal ulcer, and gastric cancer [1]. A critical step in the establishment and persistence of infection is bacterial adhesion to the gastric mucosa, which allows the organism to resist mechanical clearance and maintain close contact with host tissues. This process is mediated by outer-membrane adhesins that recognize host glycans and other surface-associated molecules, thereby shaping tissue tropism, colonization efficiency, and host–pathogen interactions [2]. Among these adhesins, the blood-group antigen-binding adhesin BabA has attracted particular attention because of its ability to recognize fucosylated blood group antigens on gastric epithelial cells and its association with persistent colonization and disease-relevant host responses [3, 4].

Targeting BabA-mediated adhesion provides a potential anti-adhesion strategy that differs from conventional antibacterial approaches aimed at killing or inhibiting bacterial growth. Such a strategy may interfere with colonization while potentially reducing selective pressure for antibiotic resistance. However, the rational development of BabA-targeting molecules requires accurate identification of functionally relevant and surface-accessible epitopes. Experimentally resolved antigen–antibody complex structures provide direct information about antibody-recognized contact regions [5], whereas computational epitope prediction can complement structural data by identifying antigenic and conserved surface regions that may not be captured by a single complex [6]. Therefore, integrating antibody-defined structural epitopes with predicted linear epitopes, conformational epitopes, and sequence conservation provides a more comprehensive basis for prioritizing candidate functional sites on BabA.

Binders, represent a promising class of engineered molecules for targeting defined protein surfaces [7]. Compared with full-length antibodies, mini-binders are smaller and structurally simpler, and they may be more amenable to computational optimization, production, and modular engineering. In particular, RFdiffusion3, a deep-learning-based generative AI model, has pushed protein design toward an all-atom design era by enabling the generation of biomolecular interaction interfaces with greater atomic-level detail [7, 8]. In this study, we sought to develop a structure-guided and epitope-prioritized computational strategy for designing mini-binders against *H. pylori* BabA. We used antibody- and nanobody-bound BabA structures to define two experimentally supported structural epitopes, and integrated BepiPred, DiscoTope, and ConSurf analyses to identify an additional candidate epitope with antigenic, conserved, and surface-exposed features. This epitope set was then used to guide the design and prioritization of compact binder candidates targeting functionally relevant surfaces of BabA.

## 2 Methods and materials

### 2.1 Structural preparation of BabA and definition of interface hotspot residues

The crystal structure of the nanobody–BabA–IgG ternary complex (PDB ID: 7ZQT) was used as the initial structural template. To define candidate hotspot residues at the BabA–binder interface, ChimeraX 1.11.1 [9] was first employed to extract BabA residues located within 4.0 Å of the binding chains, and these residues were considered candidate interface residues. The wild-type complex was then analyzed using Rosetta Interface Analyzer [10] to calculate the baseline interface energy. Subsequently, each candidate BabA interface residue was individually mutated to alanine, and Rosetta InterfaceAnalyzer was rerun for each mutant complex. The change in binding energy caused by each alanine mutation was calculated as ΔΔG_binding_ = dG_separated (mutant)_ − dG_separated (wild-type)_. Residues whose alanine substitution resulted in a marked increase in dG_separated were defined as candidate interface hotspot residues. In parallel, the BabA structure was isolated and subjected to loop refinement and structural optimization using MODELLER [11], resulting in an optimized target model for subsequent computational design.

### 2.2 Sequence- and structure-based prediction of conserved antigenic sites in BabA

BLASTP searches were performed in UniProt using UniProtKB reference proteomes + Swiss-Prot as the target database [12], with the taxonomic scope restricted to *Helicobacter pylori*. To ensure the reliability of subsequent conservation analysis, the retrieved sequences were filtered according to sequence coverage, homology confidence, sequence integrity, and redundancy. Representative sequences with high query coverage, clear homologous relationships, and origins from different strains or subtypes were retained, whereas fragments, partial sequences, low-coverage hits, sequences with abnormal lengths, and highly redundant entries were excluded. The selected homologous sequences were then subjected to multiple sequence alignment using Clustal Omega [13], and the resulting Multiple Sequence Alignment (MSA) file was exported for downstream analysis.

Evolutionary conservation analysis was performed using ConSurf [14] in PDB_MSA mode, in which the target protein PDB structure and the prepared MSA file were jointly submitted to calculate and map residue-level conservation scores onto the three-dimensional structure. This analysis was used to identify conserved regions that remain highly preserved across different *H. pylori* strains or subtypes. In parallel, BepiPred-3.0 [15] was used to predict potential linear B-cell epitopes from the primary sequence of BabA, while DiscoTope-3.0 [16] was applied to predict potential conformational B-cell epitopes based on the three-dimensional structure. The conservation results, epitope prediction outputs, and structural surface-exposure information were integrated to prioritize candidate antigenic regions. Residues or continuous segments showing high predicted antigenicity, strong evolutionary conservation, and favorable surface exposure were selected as candidate antigenic sites for subsequent experimental validation.

### 2.3 *De novo* binder backbone generation using RFdiffusion3

*De novo* protein binder backbones were generated using RFdiffusion3 [8]. Backbone generation was performed under epitope-constrained conditions, with the Refined BabA structure serving as the target and the predefined core receptor-binding epitope serving as the interface constraint. This setup was intended to bias backbone generation toward the functional binding surface of BabA and to promote the formation of candidate interfaces with favorable geometric complementarity to the target epitope. Multiple candidate binder backbones were generated for subsequent sequence design and structural evaluation.

### 2.4 Sequence design using ProteinMPNN

For each selected backbone, amino acid sequences were designed using ProteinMPNN [17]. Sequence design was carried out under structural constraints derived from the generated backbone models, with the objective of maximizing sequence–structure compatibility while preserving the intended target-binding geometry. Candidate sequences with favorable design characteristics were retained for downstream structural validation.

### 2.5 Structural back-validation of designed sequences

To evaluate whether the designed sequences were capable of folding into structures consistent with their corresponding design models, back-validation was performed using RoseTTAFold3-based structure prediction [18]. Predicted structures were compared with the original design backbones to assess structural agreement, including overall fold preservation, backbone consistency, and maintenance of the intended interface architecture. This step was used to eliminate candidates showing substantial deviation from the initial design geometry.

### 2.6 Multi-parameter prioritization of candidate binders

Candidate binders were prioritized using a composite screening framework integrating multiple structural and developability-related criteria. we employed HADDOCK 2.4 [19] for molecular docking, ProtSol [20] for solubility prediction, ProtParam [21] to estimate half-life and instability index, and PRODIGY [22] for contact-based prediction of binding affinity in protein–protein complexes. To further evaluate the dynamic stability of selected BabA–binder complexes, molecular dynamics simulations were performed using GROMACS [23]. Each complex was independently simulated twice, with each replicate run lasting 100 ns, to assess the reproducibility and stability of the predicted binding mode. The resulting trajectories were analyzed in terms of RMSD, RMSF and interfacial contact stability.

## 3 Results

### 3.1 Experimentally resolved BabA immune complexes define interface hotspot regions for binder design

To define structurally supported target residues for BabA binder design, we analyzed the experimentally resolved BabA–IgG–nanobody complex structure (PDB: 7ZQT) and identified BabA residues located within 4.0 Å of the IgG Fv or nanobody as antibody-recognized structural contact residues. These contact residues were not randomly distributed across the BabA surface but formed spatially clustered interface regions, providing structure-supported epitope patches for downstream binder design.

Rosetta computational alanine scanning was then used to estimate the energetic contribution of each antibody-recognized BabA interface residue, with positive ΔΔG_binding_ values indicating reduced interface stability after alanine substitution. Several alanine substitutions caused positive ΔΔG changes, indicating reduced predicted interface stability after mutation (Figure 1). Among them, I437A, S365A, and E444A exceeded the strong hotspot threshold of ΔΔG ≥ 2.0, suggesting that I437, S365, and E444 were the most prominent predicted interface hotspot residues. Additional residues, including L52, D233, Q207, T48, V243, and N435, exceeded the candidate hotspot threshold of ΔΔG ≥ 1.0 and were therefore retained as candidate hotspot residues. In contrast, several mutations showed low or negative ΔΔG values, suggesting limited or unfavorable energetic contribution to the modeled interface under this analysis. Therefore, the experimentally resolved BabA immune-complex structure, combined with residue-level alanine scanning, defined three groups of BabA hotspot residues for subsequent epitope prioritization and constrained *de novo* binder generation.

**Figure 1.**
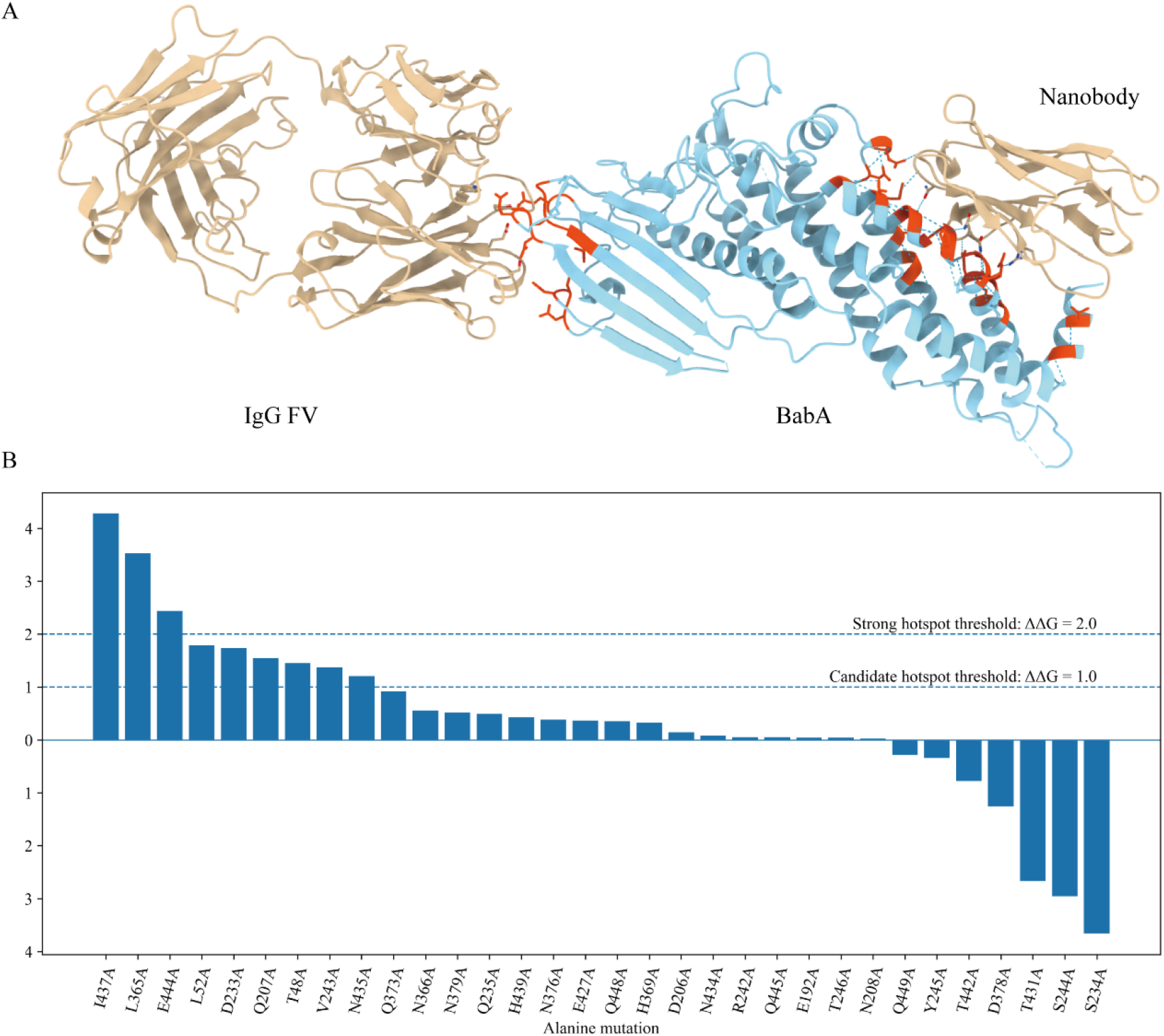
Identification and hotspot assessment of BabA interface residues in the BabA–IgG– nanobody complex. Note: A, Overall structure of the BabA–IgG–nanobody complex (PDB: 7ZQT), with BabA shown in cyan and the IgG and nanobody shown in Tan. BabA residues located within 4.0 Å of the binding chains are highlighted in orange-red. B, the hotspot residue assessment of BabA interface residues based on Rosetta computational alanine scanning.

### 3.2 Refinement generated a quality-controlled BabA structure for downstream analysis

Because several residues in the BabA structure deposited in the PDB were not experimentally resolved, structural refinement was performed to obtain a more complete and reliable model for downstream analysis (Figure 2). To obtain a BabA structure suitable for epitope mapping and binder design, the BabA chain extracted from the BabA–IgG–nanobody complex structure was subjected to MODELLER-based structural refinement. A total of ten refined models were generated and evaluated using GA341 and normalized zDOPE scores. All refined models showed GA341 values of 1.0, exceeding the commonly used reliability threshold of 0.8, indicating that the generated models were structurally reliable (Table 1). The zDOPE scores of the refined models ranged from −1.84845 to −1.76142, and all values were below 0, supporting the overall quality of the refined structures. Among them, model #2.10 showed the lowest zDOPE score of −1.84845 and was therefore selected for subsequent structural analyses.

**Figure 2.**
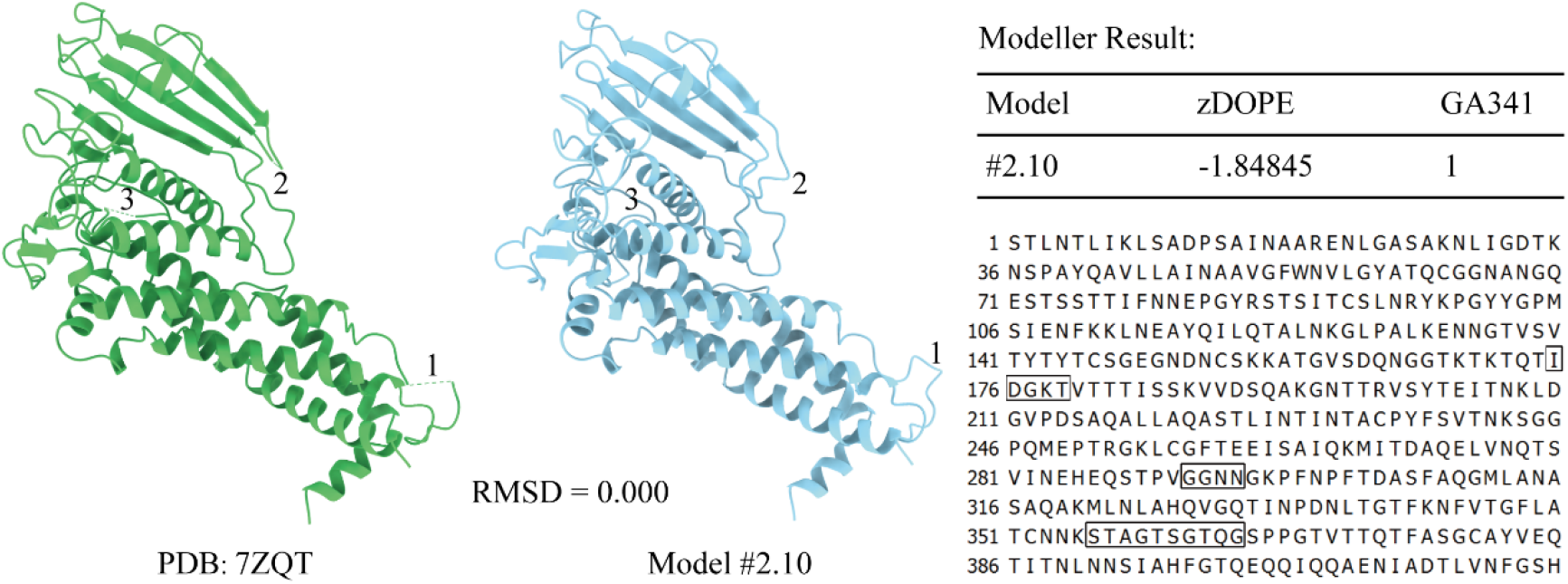
Structural refinement and quality assessment of BabA. Note: The BabA structure extracted from PDB 7ZQT is shown in green, and the MODELLER-refined BabA model #2.10 is shown in cyan. The numbered regions in the structural models and the boxed regions in the primary sequence indicate structurally inspected or refined regions during model preparation. RMSD is a statistical measure of the differences between two structures.

**Table 1.**
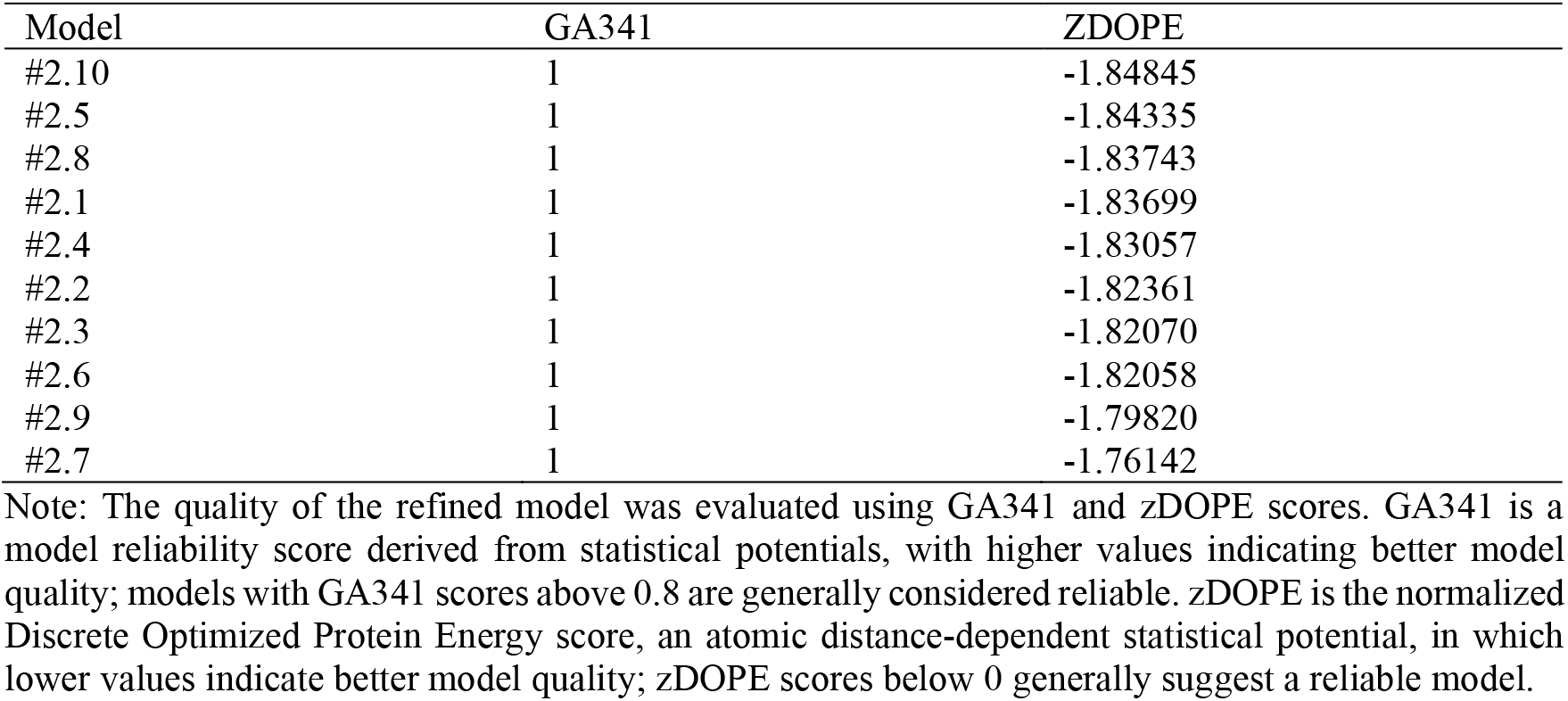
Quality assessment of MODELLER-refined BabA models.

| Model | GA341 | ZDOPE |
| --- | --- | --- |
| #2.10 | 1 | -1.84845 |
| #2.5 | 1 | -1.84335 |
| #2.8 | 1 | -1.83743 |
| #2.1 | 1 | -1.83699 |
| #2.4 | 1 | -1.83057 |
| #2.2 | 1 | -1.82361 |
| #2.3 | 1 | -1.82070 |
| #2.6 | 1 | -1.82058 |
| #2.9 | 1 | -1.79820 |
| #2.7 | 1 | -1.76142 |

Note: The quality of the refined model was evaluated using GA341 and zDOPE scores. GA341 is a model reliability score derived from statistical potentials, with higher values indicating better model quality; models with GA341 scores above 0.8 are generally considered reliable. zDOPE is the normalized Discrete Optimized Protein Energy score, an atomic distance-dependent statistical potential, in which lower values indicate better model quality; zDOPE scores below 0 generally suggest a reliable model.

Structural superposition between the BabA structure extracted from PDB 7ZQT and the selected MODELLER-refined model #2.10 showed no apparent global deviation in the displayed alignment, indicating that model #2.10 preserved the overall BabA fold while providing a quality-controlled structure suitable for downstream epitope prediction, hotspot analysis, and binder design.

### 3.3 Integrated conservation and B-cell epitope mapping prioritizes a non-redundant BabA surface region

In addition to using resources from the PDB database, we integrated evolutionary conservation analysis, surface-exposed region analysis, and linear and conformational B-cell epitope prediction to identify additional BabA regions suitable for binder design. ConSurf analysis showed that conservation across the BabA sequence was heterogeneously distributed, with both variable residues and highly conserved residues present in the structural model (Figure 3A). Residues annotated as both exposed and conserved were considered particularly important for epitope prioritization because they combined surface accessibility with evolutionary stability.

**Figure 3.**
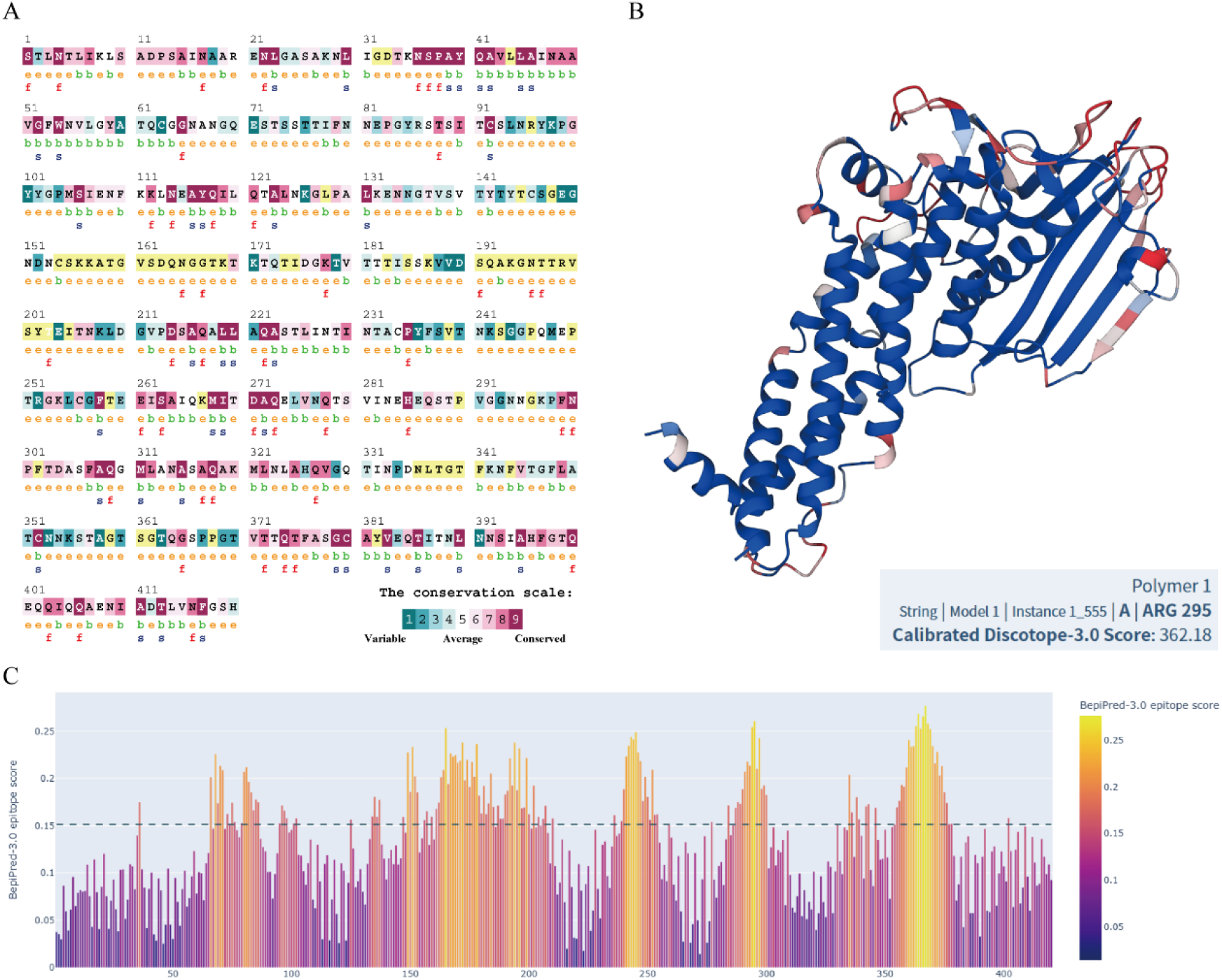
Integrated conservation analysis and B-cell epitope prediction of BabA. Note: A, Evolutionary conservation analysis of BabA using ConSurf. Different colors indicate residue conservation levels ranging from variable to conserved. The annotations below the sequence indicate predicted structural exposure and potential functional or structural relevance: e denotes an exposed residue predicted by the neural network algorithm; b denotes a buried residue; f denotes a predicted functional residue, defined as a highly conserved and exposed residue; s denotes a predicted structural residue, defined as a highly conserved and buried residue; and x denotes insufficient data, where the calculation for that site was performed on less than 10% of the sequences. B, Conformational B-cell epitope prediction of BabA using DiscoTope-3.0. Predicted conformational epitope regions were mapped onto the three-dimensional BabA structure to visualize spatially clustered antigenic regions. The DiscoTope-3.0 score increases from blue to red, indicating increasing epitope propensity. C, Linear B-cell epitope prediction of BabA using BepiPred-3.0. The bar plot shows residue-level epitope scores along the BabA primary sequence, with higher scores indicating a greater likelihood of linear B-cell epitope formation. The dashed line represents the prediction threshold, and regions above this threshold were considered potential linear B-cell epitope regions.

Structure-based conformational epitope prediction identified multiple spatially clustered antigenic regions on the BabA structure (Figure 3B). These predicted conformational epitopes were mainly located in exposed loop or surface regions rather than being uniformly distributed throughout the protein. Meanwhile, BepiPred-3.0 analysis identified multiple linear epitope-prone segments along the BabA sequence, several of which showed prediction scores above the threshold of 0.15 (Figure 3C). The major predicted linear epitope peaks were located approximately within residues 60– 90, 150–205, 240–270, 290–305, and 350–380.

To compare the predicted antigenic regions with the experimentally supported immune-recognition interface, we performed a visual analysis. Based on spatial clustering, surface exposure, and overlap with predicted epitope signals, four BabA surface regions were defined (Figure 4A). Among them, regions 3 and 4 corresponded to missing regions or missing residues. Although these regions showed high predicted antigenicity scores after MODELLER-based reconstruction, they lacked direct structural support from the experimentally resolved BabA–IgG–nanobody interface in PDB 7ZQT and were therefore not included in subsequent analyses. Region 1 was likely associated with IgG binding and potentially overlapped with the known antibody-recognition interface in the PDB structure. Therefore, hotspot residue definition in this region was primarily based on experimentally supported antibody-contact residues, with computational epitope prediction used as complementary evidence. For example, residues or regions that were predicted to be antigenic only by computational analysis but were distant from the experimentally supported immune-recognition interface in the BabA–antibody/nanobody complex were excluded from subsequent hotspot definition and binder design. In contrast, residues located close to the experimentally supported interface and showing both predicted antigenicity and high evolutionary conservation were included as candidate hotspot residues for further analysis. Region 2 retained favorable predicted epitope features and represented a non-redundant BabA surface region. This region combined sequence-level epitope propensity, conformational epitope potential, surface accessibility, and evolutionary conservation, and was therefore selected for subsequent constrained hotspot residue analysis.

**Figure 4.**
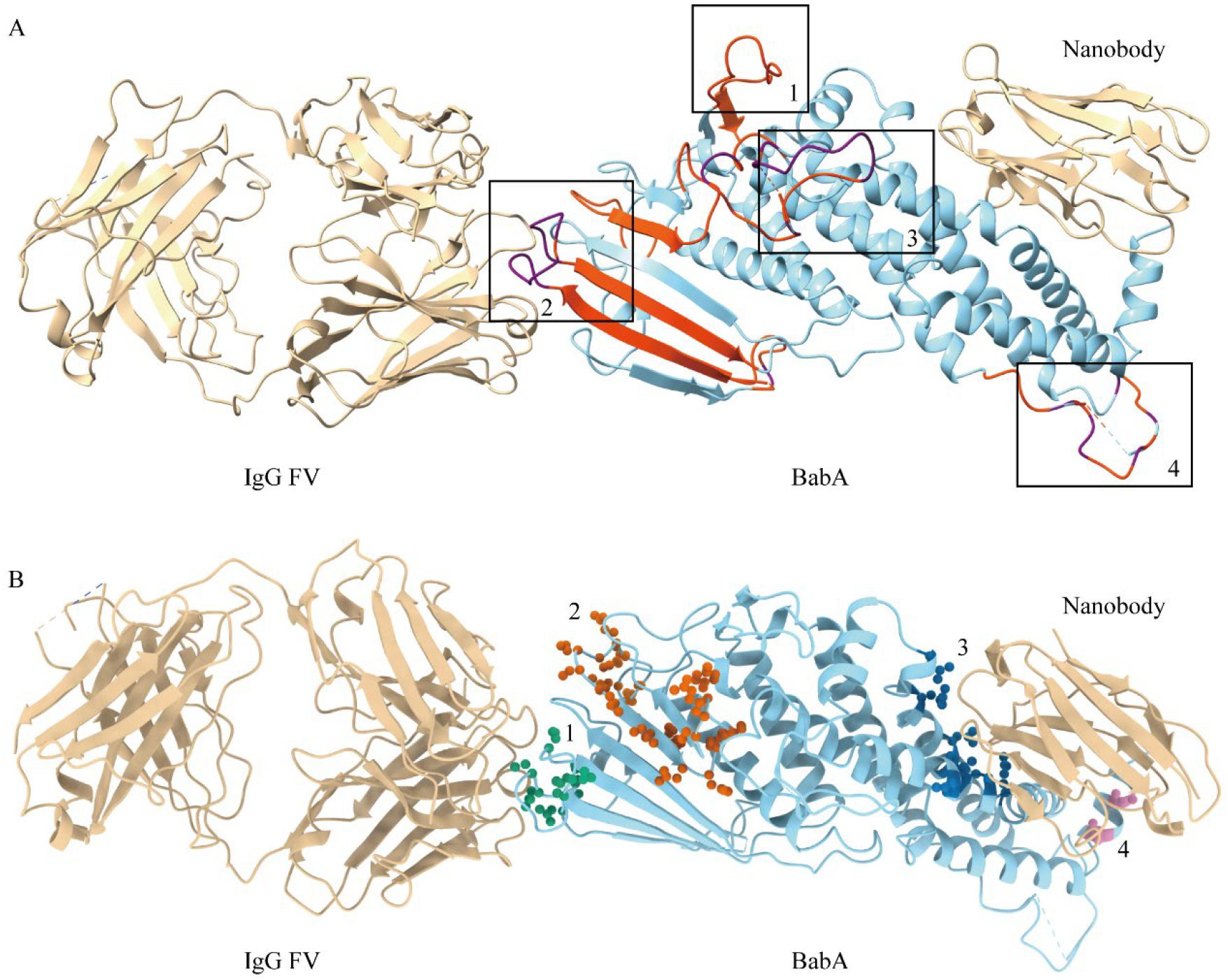
Definition and mapping of hotspot residues in the BabA–IgG–nanobody complex. Note: A, the overall structure of the BabA–IgG–nanobody complex, with BabA shown in cyan and the IgG and nanobody shown in Tan. The predicted B-cell linear and conformational epitopes are highlighted in red and purple, respectively, and the black boxes indicate candidate functional epitope regions selected based on epitope prediction, surface exposure, and interface location. B, the spatial distribution of hotspot residues selected after integrating epitope conservation into the complex structure. The four groups of candidate hotspot residues are shown as spheres in green, vermilion, dark blue, and purple, corresponding to candidate epitope regions 1–4, respectively.

**Figure 5.**
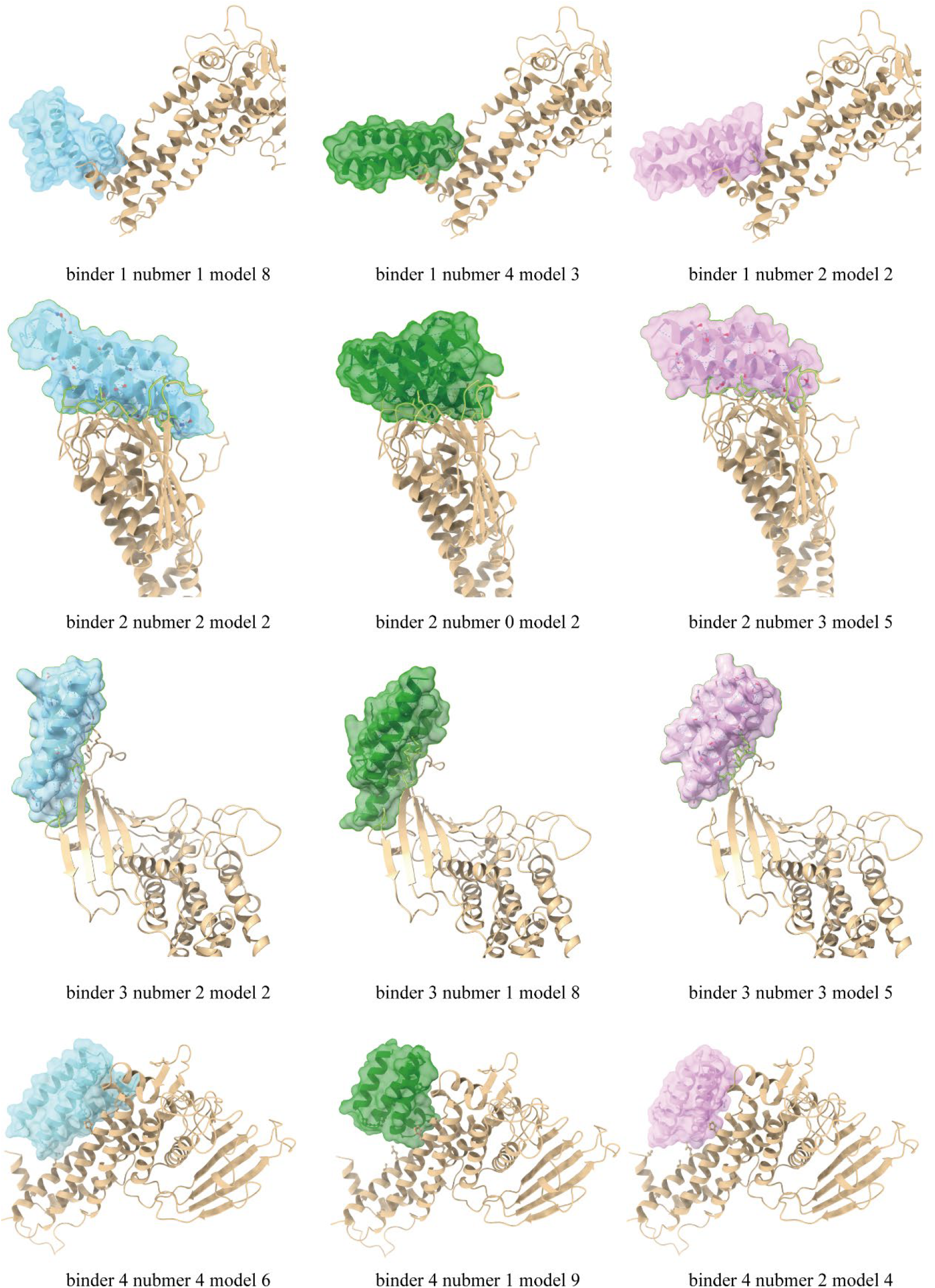
Visualization of RFdiffusion3-generated BabA–binder candidate complexes. Note: BabA is shown in tan, whereas the designed binder backbones are displayed as semi-transparent surfaces in cyan, green, or pink. “Binder 1–4” denotes the four predefined BabA hotspot or epitope regions used to guide constrained backbone generation. “Number” indicates one of the five independent RFdiffusion3 generation batches performed for each hotspot group, and “model” indicates one of the ten candidate backbone models generated in each batch. The 12 displayed candidates were retained after filtering based on structural integrity, spatial compatibility with the target epitope, compactness, secondary-structure regularity, and absence of severe clashes or extra chain breaks.

### 3.4 Constrained *de novo* backbone generation produced compact binders targeting prioritized BabA

To generate compact protein binders against the prioritized BabA epitope regions (Figure 4B), constrained *de novo* backbone generation was performed using RFdiffusion3. The selected BabA hotspot residues were used as spatial constraints to guide binder placement and interface formation. For each hotspot group, five independent design batches were performed. In each batch, a binder length was randomly sampled within the range of 65–75 amino acids, and ten backbone designs were generated based on the sampled length. In total, 200 candidate binder backbones were generated across the four hotspot groups.

The generated backbones were then filtered according to structural integrity, spatial compatibility, secondary-structure regularity, radius of gyration, and the absence of severe backbone breaks or steric clashes. After this filtering step, three candidates were retained for each hotspot group, resulting in a final set of 12 binder backbones for subsequent sequence design (Table 2). Most retained candidates adopted compact helical or mixed secondary-structure architectures and were oriented toward the predefined BabA epitope surfaces.

**Table 2.**
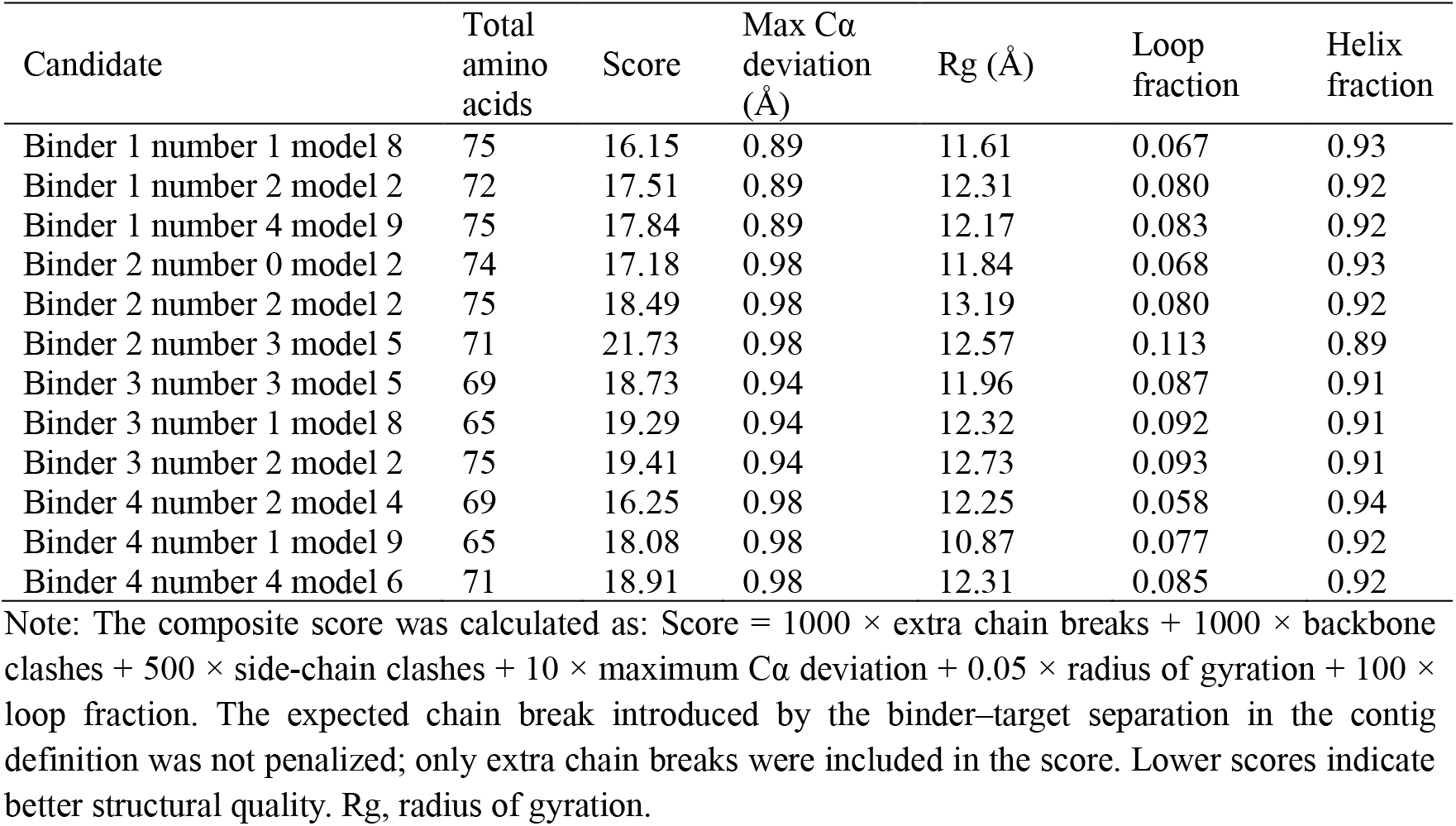
Parameters of candidate binders generated and filtered by RFdiffusion 3.

| Candidate | Total amino acids | Score | Max Cα deviation (Å) | Rg (Å) | Loop fraction | Helix fraction |
| --- | --- | --- | --- | --- | --- | --- |
| Binder 1 number 1 model 8 | 75 | 16.15 | 0.89 | 11.61 | 0.067 | 0.93 |
| Binder 1 number 2 model 2 | 72 | 17.51 | 0.89 | 12.31 | 0.080 | 0.92 |
| Binder 1 number 4 model 9 | 75 | 17.84 | 0.89 | 12.17 | 0.083 | 0.92 |
| Binder 2 number 0 model 2 | 74 | 17.18 | 0.98 | 11.84 | 0.068 | 0.93 |
| Binder 2 number 2 model 2 | 75 | 18.49 | 0.98 | 13.19 | 0.080 | 0.92 |
| Binder 2 number 3 model 5 | 71 | 21.73 | 0.98 | 12.57 | 0.113 | 0.89 |
| Binder 3 number 3 model 5 | 69 | 18.73 | 0.94 | 11.96 | 0.087 | 0.91 |
| Binder 3 number 1 model 8 | 65 | 19.29 | 0.94 | 12.32 | 0.092 | 0.91 |
| Binder 3 number 2 model 2 | 75 | 19.41 | 0.94 | 12.73 | 0.093 | 0.91 |
| Binder 4 number 2 model 4 | 69 | 16.25 | 0.98 | 12.25 | 0.058 | 0.94 |
| Binder 4 number 1 model 9 | 65 | 18.08 | 0.98 | 10.87 | 0.077 | 0.92 |
| Binder 4 number 4 model 6 | 71 | 18.91 | 0.98 | 12.31 | 0.085 | 0.92 |

Note: The composite score was calculated as: Score = 1000 × extra chain breaks + 1000 × backbone clashes + 500 × side-chain clashes + 10 × maximum Cα deviation + 0.05 × radius of gyration + 100 × loop fraction. The expected chain break introduced by the binder–target separation in the contig definition was not penalized; only extra chain breaks were included in the score. Lower scores indicate better structural quality. Rg, radius of gyration.

The retained candidates showed RFdiffusion3 scores ranging from 16.15 to 21.73, maximum Cα deviations between 0.89 and 0.98 Å, and radii of gyration ranging from 10.87 to 13.19 Å. These values indicated that the selected models remained compact and geometrically consistent with the intended backbone designs. The secondary-structure composition of the candidate binders was dominated by α-helical content, with helix fractions ranging from 0.89 to 0.94 and loop fractions ranging from 0.058 to 0.113. Among these models, Binder 1 number 1 model 8 and Binder 4 number 2 model 4 showed the lowest RFdiffusion3 scores, 16.15 and 16.25, respectively, while maintaining low Cα deviation and high helical content.

Taken together, constrained *de novo* backbone generation produced a focused set of structurally plausible and compact BabA-targeting binder backbones with favorable geometric features. These candidates were therefore advanced to amino acid sequence design, structural back-validation, and BabA–binder complex evaluation.

### 3.5 Sequence design and structural back-validation refined candidate BabA binders

After ProteinMPNN-based sequence design and RoseTTAFold3-based structural back-validation, one representative sequence was retained for each of the 12 RFdiffusion3-generated binder backbones according to sequence–backbone compatibility, structural agreement, fold compactness, and BabA-facing interface geometry. The retained candidates showed ProteinMPNN scores ranging from 0.7407 to 1.0189, with most candidates displaying low structural deviation from their corresponding design backbones after back-validation (Table 3). In particular, Binder 4191, Binder 2223, Binder 1493, Binder 2351, Binder 3352, and Binder 3182 showed RMSD values below 0.31 Å, indicating strong agreement between the designed backbone and the independently predicted structure. Binder 4461 also maintained an acceptable RMSD of 0.686 Å, whereas Binder 3221 showed a markedly higher RMSD of 4.525 Å, suggesting a substantial deviation from the intended backbone despite its favorable ProteinMPNN score. These results indicate that most designed sequences were able to preserve the intended compact fold and target-facing geometry, while candidates with larger structural deviations were treated with caution in downstream prioritization.

**Table 3.** ProteinMPNN-based sequence design and RoseTTAFold3 structural back-validation of candidate BabA binders.

| Name | Sample | Score | Seq recovery | RMSD (Å) |
| --- | --- | --- | --- | --- |
| Binder 1181 | Binder 1 number 1 model 8 rank 1 | 0.8893 | 0.3733 | 0.418 |
| Binder 1225 | Binder 1 number 2 model 2 rank 5 | 1.0189 | 0.2917 | 0.345 |
| Binder 1493 | Binder 1 number 4 model 9 rank 3 | 0.7407 | 0.3467 | 0.285 |
| Binder 2024 | Binder 2 number 0 model 2 rank 4 | 0.9534 | 0.4730 | 0.434 |
| Binder 2223 | Binder 2 number 2 model 2 rank 3 | 0.9178 | 0.4000 | 0.276 |
| Binder 2351 | Binder 2 number 3 model 5 rank 1 | 0.8922 | 0.4225 | 0.298 |
| Binder 3221 | Binder 3 number 2 model 2 rank 1 | 0.8825 | 0.3692 | 4.525 |
| Binder 3352 | Binder 3 number 3 model 5 rank 2 | 0.8866 | 0.4058 | 0.299 |
| Binder 3182 | Binder 3 number 1 model 8 rank 2 | 0.8997 | 0.3385 | 0.307 |
| Binder 4191 | Binder 4 number 1 model 9 rank 1 | 0.7882 | 0.5385 | 0.225 |
| Binder 4244 | Binder 4 number 2 model 4 rank 4 | 0.9309 | 0.4493 | 0.409 |
| Binder 4461 | Binder 4 number 4 model 6 rank 1 | 0.9645 | 0.4085 | 0.686 |

Note: For each RFdiffusion3-generated binder backbone, 64 sequences were designed using ProteinMPNN. The top five sequences with the lowest ProteinMPNN scores were subjected to RoseTTAFold3-based structural back-validation. One sequence showing the best overall structural agreement, folding confidence, compactness, and BabA-facing interface geometry was retained for each backbone.

**Table 4.**
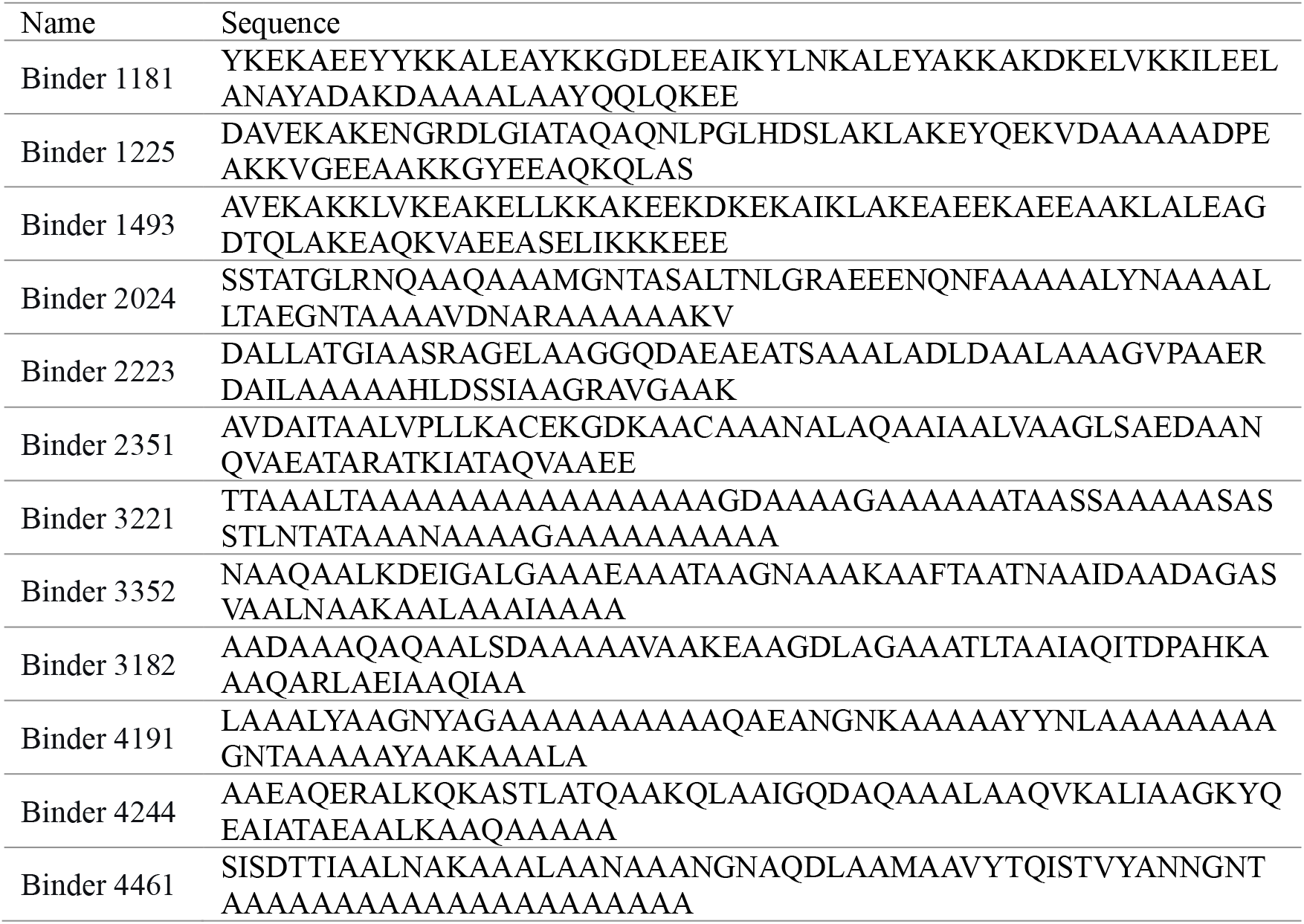
Amino acid sequences of designed binders.

Together, this sequence design and back-validation step reduced the candidate pool to 12 sequence-defined BabA binder models with varying levels of structural confidence. These candidates were subsequently advanced to molecular docking, interface energy evaluation, and dynamic stability analysis.

### 2.5 Binding modes and stability of BabA–binder complexes

To evaluate whether the selected binders could effectively bind to the prioritized surface region of BabA, molecular docking was first performed between BabA and the 12 structurally validated candidate binders (Figure 6). The docking models were selected by comprehensively considering the internal HADDOCK scoring parameters, the spatial occupancy of the predefined BabA epitope region by each candidate binder, shape complementarity, and residue-level contact patterns. The results showed that different binders adopted distinct binding orientations on the BabA surface. Some candidates covered the predefined functional epitope region and formed relatively compact interfacial contacts, suggesting that these designed binders may have the potential to target the functional surface of BabA.

Based on these docking results, we further integrated the predicted binding free energy, dissociation constant, solubility, half-life, and instability index to comprehensively evaluate the developability potential of the 12 pre-selected binders (Table 5). All candidate binders displayed varying degrees of predicted binding capacity, with ΔG values ranging from –8.1 to –14.5 kcal·mol^−1^ and corresponding K_d_ values ranging from 1.1 × 10^−6^ to 2.4 × 10^−11^ M. Among them, Binder 4461 showed the strongest predicted binding affinity, with a ΔG of –14.5 kcal·mol^−1^ and a K_d_ of 2.4 × 10^−11^ M. It also exhibited a high predicted solubility score of 0.9825 and a low instability index of 3.33, suggesting a favorable combination of strong binding potential, good solubility, and high structural stability. Binder 3352 also showed excellent overall properties, with a ΔG of –13.7 kcal·mol^−1^, a K_d_ of 8.3 × 10^−11^ M, a solubility score of 0.9834, and an instability index of 3.09. Binder 2024, Binder 2351, and Binder 3182 likewise displayed predicted nanomolar-level binding affinity and relatively low instability indices, and therefore may serve as backup candidates for subsequent validation. In contrast, although Binder 3221 showed strong predicted binding ability, with a ΔG of –12.2 kcal·mol^−1^ and a K_d_ of 1.1 × 10^−9^ M, it was predicted to be insoluble, indicating a potentially high risk during subsequent expression and purification. Binder 1181 showed relatively weak predicted affinity, with a ΔG of –8.1 kcal·mol^−1^ and a K_d_ of 1.1 × 10^−6^ M, and was classified as unstable; therefore, it was assigned a lower priority.

**Table 5.** physicochemical properties of pre-selected binders.

| Name | $\Delta G$ (kcal mol<sup>-1</sup>) | K<sub>d</sub> (M) | Solubility | Half-life | Instability index |
| --- | --- | --- | --- | --- | --- |
| Binder 1181 | -8.1 | 1.1e-06 | 0.9975 | 2.8 hours<sup>1</sup>; 2 min<sup>2</sup> | unstable |
| Binder 1225 | -10.0 | 4.8e-08 | 0.9839 | 1.1 hours<sup>1</sup>; >10 hours<sup>2</sup> | 16.94 |
| Binder 1493 | -9.2 | 1.7e-07 | 0.9978 | 4.4 hours<sup>1</sup>; >10 hours<sup>2</sup> | 26.07 |
| Binder 2024 | -12.1 | 1.4e-09 | 0.9570 | 1.9 hours<sup>1</sup>; >10 hours<sup>2</sup> | 10.53 |
| Binder 2223 | -10.8 | 1.1e-08 | 0.9541 | 1.1 hours<sup>1</sup>; >10 hours<sup>2</sup> | 21.69 |
| Binder 2351 | -11.3 | 4.9e-09 | 0.9473 | 4.4 hours<sup>1</sup>; >10 hours<sup>2</sup> | 22.17 |
| Binder 3221 | -12.2 | 1.1e-09 | Insoluble | 7.2 hours<sup>1</sup>; >10 hours<sup>2</sup> | 11.61 |
| Binder 3352 | -13.7 | 8.3e-11 | 0.9834 | 1.4 hours<sup>1</sup>; >10 hours<sup>2</sup> | 3.09 |
| Binder 3182 | -11.6 | 3.1e-09 | 0.9466 | 4.4 hours<sup>1</sup>; >10 hours<sup>2</sup> | 15.50 |
| Binder 4191 | -11.7 | 2.4e-09 | 0.9938 | 5.5 hours<sup>1</sup>; 2 min<sup>2</sup> | 17.47 |
| Binder 4244 | -9.9 | 5.4e-08 | 0.9902 | 4.4 hours<sup>1</sup>; >10 hours<sup>2</sup> | 18.80 |
| Binder 4461 | -14.5 | 2.4e-11 | 0.9825 | 1.9 hours<sup>1</sup>; >10 hours<sup>2</sup> | 3.33 |

Note: $\Delta G$ is the predicted binding free energy; K<sub>d</sub> values are calculated at 25.0 °C. Solubility is predicted using a model with a threshold of 0.84, which minimizes false positives (i.e., rarely misclassifies insoluble proteins as soluble) at the cost of potentially discarding some soluble proteins – suitable for screening aimed at experimental purification. Superscript <sup>1</sup> indicates half-life in mammalian reticulocytes, which simulates the eukaryotic degradation machinery and is commonly used to predict the therapeutic window of a target protein in mammals or humans; superscript <sup>2</sup> indicates half-life in *E. coli*, reflecting the protein's ability to resist degradation by the host's endogenous proteases. Instability index: values <40 indicate stable proteins; values >40 indicate unstable proteins.

To further evaluate the dynamic stability of the prioritized BabA–binder complex, Binder 3352 was selected for molecular dynamics simulation because it showed the most favorable docking performance among the candidate binders, with a predicted binding free energy of −13.7 kcal mol^−1^ and an estimated Kd of 8.3 × 10^−11^ M (Figure 7A). During two independent 100 ns simulations, the backbone RMSD of the BabA–Binder 3352 complex increased rapidly during the initial relaxation phase and subsequently reached a relatively stable plateau (Figure 7B). In run 1, the RMSD rose from approximately 1.0 Å to around 2.0 Å within the first 10 ns, followed by moderate fluctuations mainly between 2.5 and 4.0 Å after approximately 25–30 ns. In run 2, the RMSD followed a comparable trend but remained slightly lower and more compact, stabilizing mostly within the range of 2.5–3.3 Å after the initial equilibration period. Importantly, neither trajectory showed a continuous increase in RMSD or abrupt structural deviation over the 100 ns simulation, suggesting that the complex did not undergo global destabilization or binder dissociation from the BabA surface.

**Figure 7.**
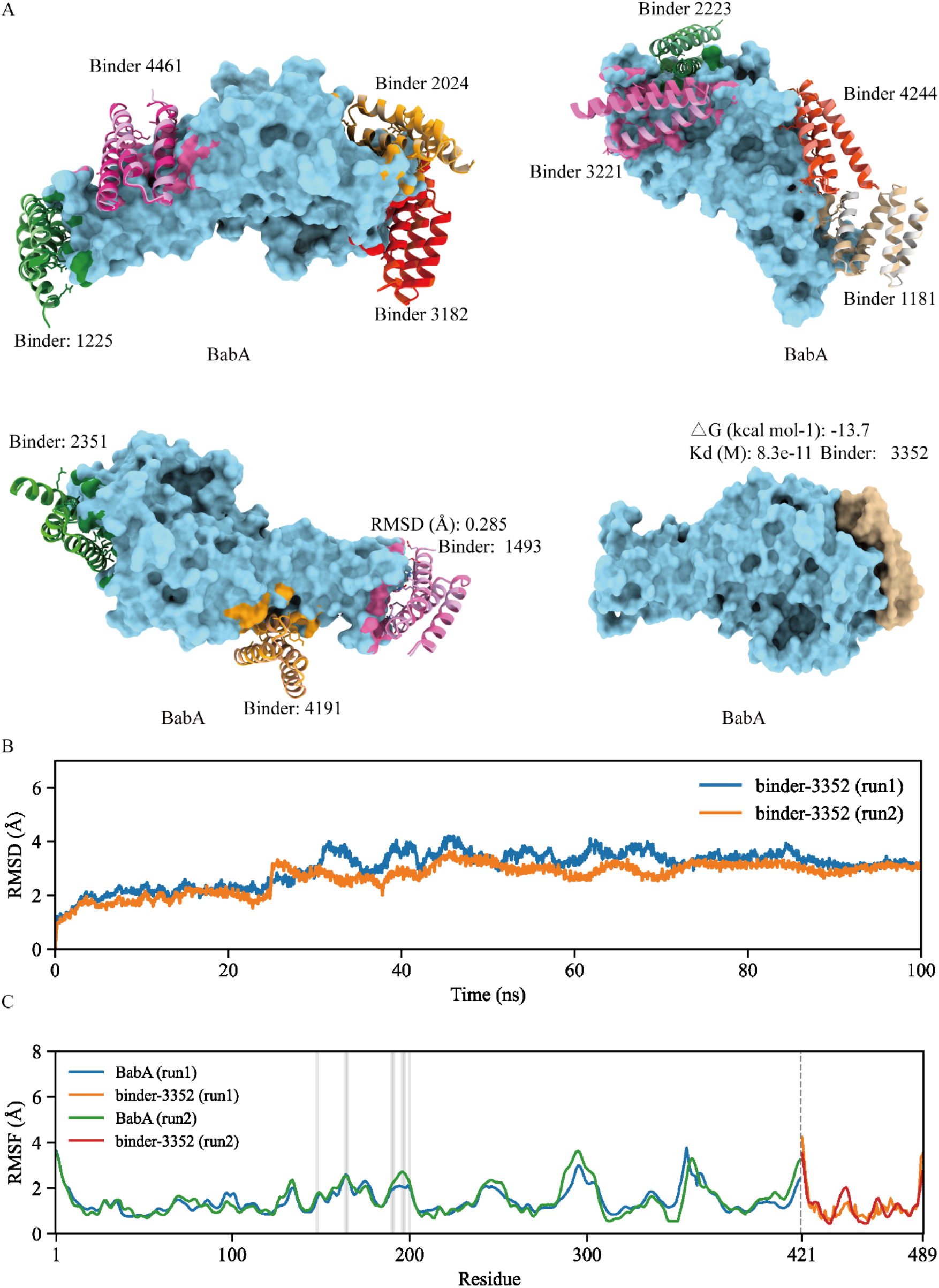
Docking models and molecular dynamics simulation of candidate BabA-binding proteins. Note: A, Molecular docking models of 12 candidate binders bound to BabA. BabA is displayed as a cyan surface, and the binders are shown in different colors. B, Backbone RMSD profiles of the BabA–Binder 3352 complex from two independent 100 ns molecular dynamics simulations. C, RMSF per residue for BabA and the binder-3352 within the complexes. BabA and binder-3352 were labeled with different colors. Grey shaded rectangles indicate the predefined hotspot residues on BabA located at the binder– interface region.

Per-residue RMSF analysis further supported the structural stability of the BabA–Binder 3352 complex (Figure 7C). For BabA, most residues displayed relatively low to moderate fluctuations, generally within approximately 1–2 Å, while higher RMSF peaks were mainly observed in several flexible regions, including the N-terminal region, residues around 280–310, residues around 350–370, and the region close to the BabA C-terminus. Notably, the predefined hotspot residues located at the binder–BabA interface, indicated by grey shaded rectangles, showed only moderate fluctuations and did not correspond to the most flexible regions of BabA. This suggests that the key interface region remained relatively constrained during the simulations. For Binder 3352, the internal residues also showed generally low RMSF values, whereas higher fluctuations were mainly observed near the terminal or boundary regions, particularly around residue 421 and the C-terminal end. Together, the consistent RMSD plateau and the restrained RMSF pattern at the interface indicate that Binder 3352 maintained a relatively stable binding conformation on the BabA surface during the simulations, supporting its potential as a dynamically stable BabA-binding candidate, supporting Binder 3352 as a dynamically stable candidate for further experimental validation.

## Discussion

*Helicobacter pylori* is an important gastrointestinal pathogen closely associated with chronic gastritis, gastric ulcer, duodenal ulcer, and gastric cancer, and has been classified as a Group I carcinogen by the International Agency for Research on Cancer under the World Health Organization [24]. Among its virulence-associated factors, BabA is a key outer-membrane adhesin that mediates host recognition and adhesion to gastric epithelial cells, thereby contributing to tissue tropism, long-term colonization, and disease-related host responses [25]. Therefore, targeting the functional surface of BabA may provide a potential anti-adhesion strategy. Experimentally resolved BabA– antibody and BabA–nanobody complex structures provide valuable information on immune-recognized functional surfaces of BabA, while sequence- and structure-based epitope prediction can further identify conserved, highly antigenic, and surface-accessible regions. However, these structural and immunological resources are usually used for epitope annotation, vaccine design, or antibody characterization, and are less frequently converted into spatial constraints for the *de novo* design of compact protein binders.

In this study, we established an epitope-guided computational design workflow that integrates three experimentally resolved structural interfaces with one computationally predicted epitope, converting them together into an effective starting point for protein binder design. This combined evidence is important because a BabA binder with application potential should not only recognize a spatially accessible surface region, but should also preferentially target regions that are functionally relevant and relatively conserved across different *H. pylori* strains. On this basis, we generated 200 candidate binder backbones around four groups of hotspot regions and selected 12 structurally reasonable candidates. ProteinMPNN was then used to generate 64 sequences for each candidate backbone, resulting in a total of 768 designed sequences. These sequences were further subjected to RoseTTAFold3-based structural back-validation, ultimately yielding 12 candidate BabA binder models. Overall, the retained candidate models were small proteins dominated by α-helical or mixed secondary-structure architectures, consistent with the design goals of structural simplicity and engineering flexibility for compact protein binders.

The multi-parameter evaluation further indicated that predicted binding strength alone is insufficient as the sole criterion for candidate selection. Binder 4461 showed the strongest predicted affinity, while also exhibiting favorable predicted solubility and a low instability index, making it one of the most promising candidates from a developability perspective. Binder 3352 also displayed strong predicted binding capacity, favorable predicted solubility, and a low instability index, and its BabA complex remained relatively stable in two independent molecular dynamics simulations. This suggests that Binder 3352 may represent a balanced candidate in terms of predicted affinity and dynamic stability. In contrast, although Binder 3221 showed favorable predicted affinity, it was predicted to be insoluble; Binder 1181 showed weaker predicted affinity and was classified as unstable. These differences indicate that prioritizing compact protein binder candidates should involve the integrated consideration of affinity-related metrics, solubility, structural stability, and dynamic behavior, rather than relying solely on predicted binding energy.

An important conceptual contribution of this study is the conversion of epitope discovery from a descriptive endpoint into a generative design input. In traditional vaccine or antibody studies, antigen epitope identification is usually regarded as an endpoint, mainly used to explain antibody-binding sites, antigen immunogenicity, or candidate vaccine regions [26, 27]. In this study, however, epitope information was further transformed into the starting point for *de novo* binder design. This shifts the research logic from “identifying where antibodies bind” to “designing new binding molecules using antibody-recognized and conserved regions.” From this perspective, existing structural databases and immunological data are no longer merely static annotations, but are instead transformed into spatially defined design constraints that can be used for AI-assisted protein engineering. Therefore, public antigen–antibody complex structures, epitope prediction results, and conservation data can be repurposed for computational protein engineering. This framework may also be transferable to other pathogen adhesins and virulence factors for which structural or epitope information is available.

Despite these advantages, several limitations should be acknowledged. First, all candidate binders in this study remain computational predictions. Molecular docking scores, predicted binding free energies, PRODIGY-derived K_d_ values, solubility scores, instability indices, and molecular dynamics simulations can be used for candidate prioritization, but they cannot replace experimental validation. Therefore, the predicted K_d_ values in this study should be interpreted as relative ranking indicators rather than definitive binding constants. Second, only Binder 3352 was subjected to molecular dynamics simulation. Although this result supports a certain degree of dynamic stability for this complex, parallel simulations of Binder 4461 and other top-ranked candidates are still needed for a more comprehensive comparison of interface stability. Third, some candidates still showed potential developability risks. For example, Binder 3221 showed favorable predicted affinity but was predicted to be insoluble, whereas Binder 1181 was classified as unstable. These results indicate that candidate selection should not rely solely on binding strength, but should also consider solubility, expression feasibility, structural stability, and dynamic interface stability. Fourth, the current workflow does not include experimental directed evolution. In many binder development pipelines, computational design usually provides only initial hits, which still require further improvement through yeast display, phage display, bacterial display, site-saturation mutagenesis, or deep mutational scanning to enhance affinity, specificity, and expression performance [28, 29]. Therefore, the candidates obtained in this study should be regarded as first-generation computational candidates rather than experimentally optimized final molecules.

Future studies should focus on experimental validation and iterative optimization. First, top-ranked candidates such as Binder 3352 and Binder 4461 should be prioritized for expression and purification to evaluate their expression levels,solubility, and production feasibility. Subsequently, their direct binding to recombinant BabA can be tested using ELISA, biolayer interferometry, surface plasmon resonance, pull-down assays, or other quantitative binding methods. Further functional experiments should assess whether these binders can block the interaction between BabA and host glycan receptors or gastric epithelial cell models. Given the sequence variation of BabA among different *H. pylori* strains, cross-strain BabA binding assays will also be needed to determine whether the conserved epitope-guided design strategy supports broader recognition. In later stages, the best-performing candidates may be further improved through interface redesign, multivalent fusion, directed evolution, or stability engineering to enhance their affinity, specificity, expression performance, and functional activity. In addition, mirror-image D-protein design may serve as a potential extension to improve resistance to proteolytic degradation and stability in the complex gastrointestinal environment [30-32].

In summary, this study proposes a transferable computational framework for converting antibody-defined and computationally prioritized epitopes into AI-assisted *de novo* binder design constraints. By integrating structural immunology, epitope prediction, conservation analysis, RFdiffusion3-based backbone generation, ProteinMPNN-based sequence design, structural back-validation, molecular docking, and molecular dynamics simulation, this study obtained a set of BabA-binding candidates with potential development value. Although experimental validation remains essential, our results suggest that epitope guidance, database resource reuse, and AI-assisted protein design can jointly provide a promising strategy for developing compact binders targeting pathogen adhesins and other important microbial surface proteins.

## Data and model availability statement

The dataset and candidate binder models generated in this study are available in the GitHub repository: https://github.com/ZhuYaojun1

## Authors’ contributions

YZ: Conceptualization, Methodology, Investigation, Formal analysis, Visualization, Writing – original draft. MBI: Writing – review & editing. XZ: Conceptualization, Supervision, Funding acquisition, Project administration, Writing – review & editing. All authors read and approved the final manuscript.

